# Matching in the wild: promise and pitfalls of propensity score matching for field ecology

**DOI:** 10.64898/2026.07.31.741844

**Authors:** Olivia Ross, Katherine Siegel, Kathy Baylis, Sara Goeking, Joan Dudney

## Abstract

Field ecologists often rely on observational data to understand the impact of environmental stressors and management interventions on natural systems. Natural and anthropogenic events (e.g. wildfires, protected areas, nutrient deposition) do not occur randomly in space, however, which can introduce bias into observational studies—which we refer to as causal selection bias. Field study designs that ignore the non-random occurrence of stressors may yield biased estimates of stressor effects on ecosystems. Matching methods commonly used in economics, political science and epidemiology offer a powerful framework for controlling for causal selection bias by identifying more comparable treatment and control sites. Although these methods are increasingly used in conservation, they are rarely used in ecological field-based studies. Here we review how Propensity Score Matching (PSM) can improve field sampling designs in ecology and strengthen causal identification of stressor effects. Then we apply this approach to a case study examining wildfire effects on forest recovery in California. We conclude with practical recommendations for implementing PSM to improve causal identification of ecological change, which is particularly important for developing effective management interventions.

## Introduction

A central goal in ecology is to understand how anthropogenic and natural disturbances shape the structure and function of ecosystems. Because manipulative experiments are often infeasible (Sagarin & Pauchard, 2009), field ecologists frequently rely on observational data collected using two classes of research designs: 1) random sampling designs (e.g. simple, stratified, or systematic) and 2) natural experiments that compare **treatment** and **control** observational units (De La Palma et al., 2018; Millstein, 2019). To understand restoration treatment efficacy, for instance, researchers might compare species richness between restored and unrestored areas. Though their implicit goal is causal understanding, these designs can overlook important sources of **statistical bias**, leading to misestimation of the effects of disturbances (Simler-Williamson & Germino, 2022; Honey & Roses et al., 2011; Andam et al., 2008). Given the rapid increase in anthropogenic-driven disturbances across ecosystems worldwide (Potapov et al., 2025; Allen et al., 2015), research designs that control for statistical bias and strengthen causal inference are critical to inform effective conservation and management interventions (Baylis et al, 2026; Jones & Shreedhar, 2024).

An important source of statistical bias arises when disturbances (anthropogenic or natural) do not occur randomly. Ecosystem characteristics that increase the probability of a disturbance or management intervention—e.g. distance to road or shoreline, topography, disturbance history— can vary systematically from the undisturbed ecosystem. Wildfires, for instance, are more likely to ignite in hotter, drier regions closer to human development (Rogeau & Armstrong, 2017) and restoration efforts are often selected based on a pre-defined criterion (e.g. accessibility, protected areas) (Chambers et al., 2019). This can lead to **causal selection bias—**when the environmental driver of interest is not randomly distributed across the sample (*Figure 1C–D*; Schafer & Kang, 2008; Ramsey et al, 2019). Field studies that do not address causal selection bias may capture effects of other drivers in the ecosystem that are correlated with the probability of the disturbance, leading to misestimation of the focal disturbance effect.

**Figure 1.**
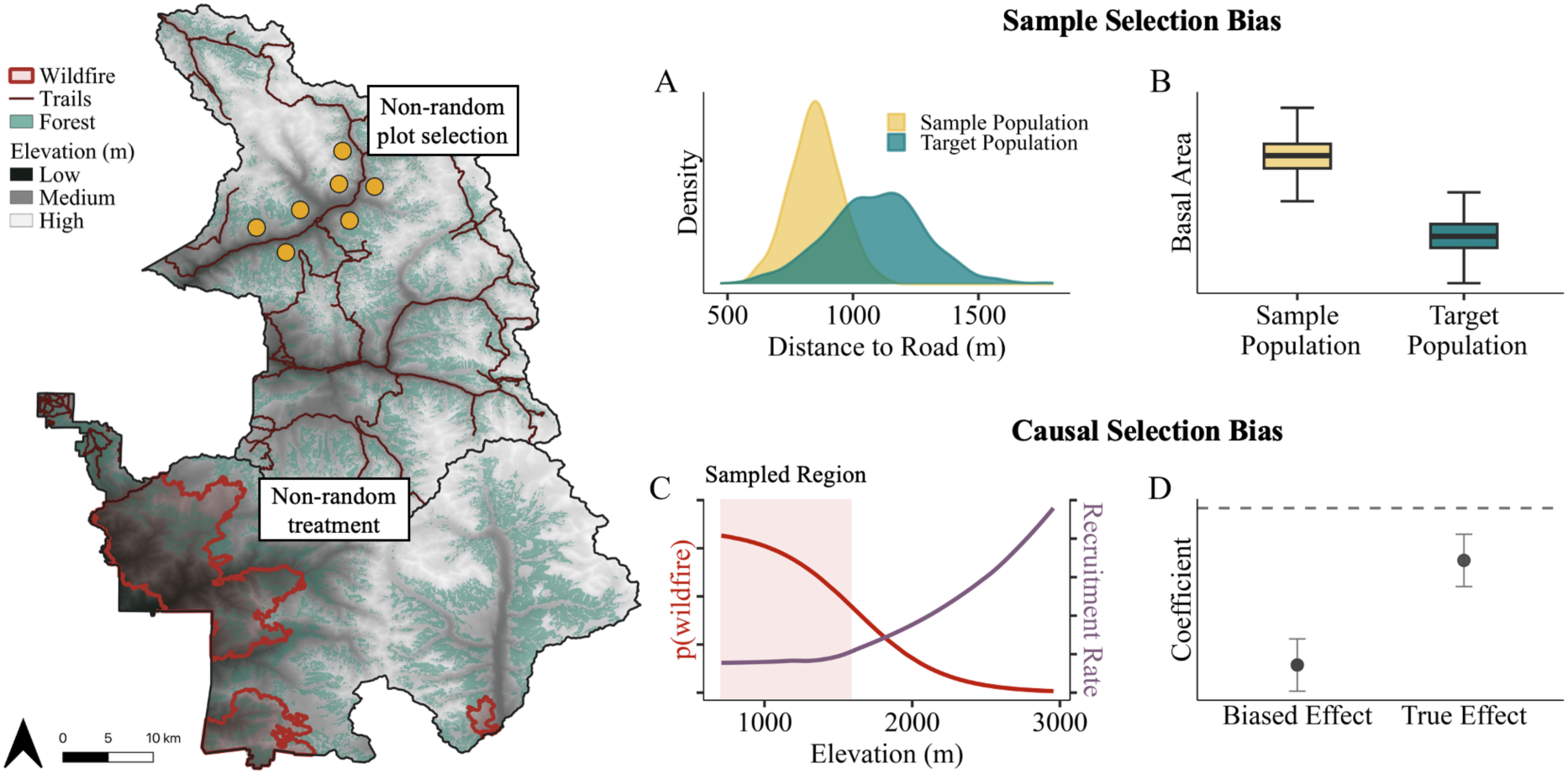
Comparison between sample selection bias (A–B) and causal selection bias (C–D). Sample selection bias occurs when the sample is not representative of the target population (A). Consequently, sample-based estimates (i.e. means or proportions) systematically deviate from their true population values (B). For example, researchers seeking to quantify forest basal area may randomly select plots that are more accessible and safer to sample (“sample population”; yellow points on the map). If roads and trails are close to waterways and deeper soils, sampled forest stands may be more productive, leading to an overestimate of the target population’s basal area (B). In contrast, causal selection bias occurs when the treatment (e.g. disturbance, management intervention) is not randomly distributed across the target sample (e.g. mixed conifer forests) (C). This non-random assignment of treatment can lead to systematic differences between treatment and control observational units that result in a biased estimate of the treatment effect (D). For example, wildfires do not occur randomly—they are more likely to occur in hotter, drier regions, climatic conditions that also independently affect tree recruitment (red polygons on the map). Unless climate is controlled for in the study design, the estimated effect of fire on forest recovery may be biased (D).

Matching methods, commonly used in economics, political science and epidemiology (Brookhart et al., 2006; Imbens, 2004), provide a critical advance for observational designs in ecology because they explicitly address causal selection bias (Rosenbaum & Rubin, 1983). By statistically identifying treatment (disturbed) and control (undisturbed) groups that experience similar conditions, matching methods improve causal identification of estimated effects (Rosenbaum & Rubin, 1983). Although matching methods have been increasingly used in ecology and conservation (Baylis et al. 2026; Schleicher et al., 2020; Siegel et al., 2022a; Siegel et al., 2022b; Woo et al., 2021a), they are rarely applied to field-based studies, where they can improve the causal identification of disturbance impacts on ecosystems. Here we outline key strengths and limitations of matching—specifically Propensity Score Matching (PSM)—in a field ecology setting. Then, we apply PSM to a case study examining the effects of wildfire on forest recovery in California. We conclude with practical considerations when using this approach and highlight how PSM can improve estimation of ecological change.

## What is Propensity Score Matching?

Propensity Score Matching is a statistical technique that identifies comparable treatment and control units (e.g. individuals, plots, sites) when the treatment is not randomized (Rosenbaum & Rubin, 1983). By pairing treatment and control units with similar characteristics, PSM enables observational studies to approximate an experimental design (referred to as a "quasi-experimental” design) (Austin, 2011a). If matched sites are comparable and assumptions of PSM are satisfied (see below), the treatment effect is causally identifiable (Rosenbaum & Rubin, 1983). Thus, PSM is a powerful tool for field-based ecological studies when experimental manipulation of treatments (e.g. restoration, fire, or hurricanes) is logistically, financially, or ethically infeasible.

Comparability between treatment and control units is achieved by matching them on variables that would otherwise lead to systematic differences between groups. These variables—referred to as **confounding variables**—are site characteristics that influence both the outcome of interest (e.g. native species recovery) and the likelihood of receiving a treatment (e.g. restoration interventions). If confounding variables are not controlled for in the study design, they can result in biased **estimated effects**, where the effect deviates (i.e. has a different sign or magnitude) from the true effect (*Figure 1D*; Dudney et al., 2025; Byrnes & Dee, 2025). After confounding variables have been identified and collated, researchers estimate a **propensity score,** which is a unitless scaler that estimates the probability that a plot experiences the treatment (Rosenbaum & Rubin, 1985). PSM statistically identifies comparable treatment and control units with the most similar propensity scores—i.e. units with similar values of all confounding variables included in the design (Rosenbaum & Rubin, 1983). The resulting matched dataset mimics a randomized experiment (Ferraro & Haunar, 2014), where treatment and control groups differ only in exposure to the treatment, if all assumptions are met (Rosenbaum & Rubin, 1983).

## Strengths of PSM applications in field ecology

### Makes the assumptions required for causal inference explicit and testable

Assumptions required for causal inference are not often explicit and testable in classical field study designs, leading to a lack of clarity as to when cause-and-effect questions can be answered. For example, the probability of treatment assignment is seldom addressed before collecting field data (Stuart, 2010), and regression-based analyses do not include standard diagnostics that assess the degree of overlap between the treatment and control groups (Rubin, 2001; Deheja and Wahba, 1999). If these two assumptions are not addressed, analyses may interpolate between two populations that differ ecologically (Austin, 2011; Stuart 2010), and identifying the treatment effect becomes infeasible.

To identify causal effects, PSM rests on three assumptions: 1) all confounders have been measured and included in the propensity score model (Stuart, 2010; Rosenbaum & Rubin, 1983), 2) each selected plot has a nonzero probability of receiving the treatment (i.e. common support condition) (Rubin, 1980), and 3) the **stable unit treatment value assumption (SUTVA)** is not violated— i.e. one unit’s treatment (e.g. wildfire) does not affect another unit’s outcome and treatments are consistently applied across units (Rubin, 1980). Each assumption can be explicitly addressed at the design stage, before field data collection, when the study can still be refined to support causal inference.

Although the first assumption is the most consequential and least testable, developing a Directed Acyclic Graph (DAG) to explicitly identify all confounding variables in a study (Arif & MacNeil, 2023) can improve transparency and allow future researchers to evaluate and refine the research design (Siegel & Dee 2025). Additionally, if confounders are known but unmeasured, sensitivity analyses can quantify the degree to which this variable might influence the treatment effect (see Considerations below) (Cinelli & Hazlett 2020; Rosenbaum, 1987; Rosenbaum, 2002). The common support condition can also be tested visually (*Figure 2*) or through statistical tests to quantify the similarity between both groups (Stuart, 2010). Finally, compliance with SUTVA can be evaluated and improved during the field design stage (Stuart, 2010).

**Figure 2.**
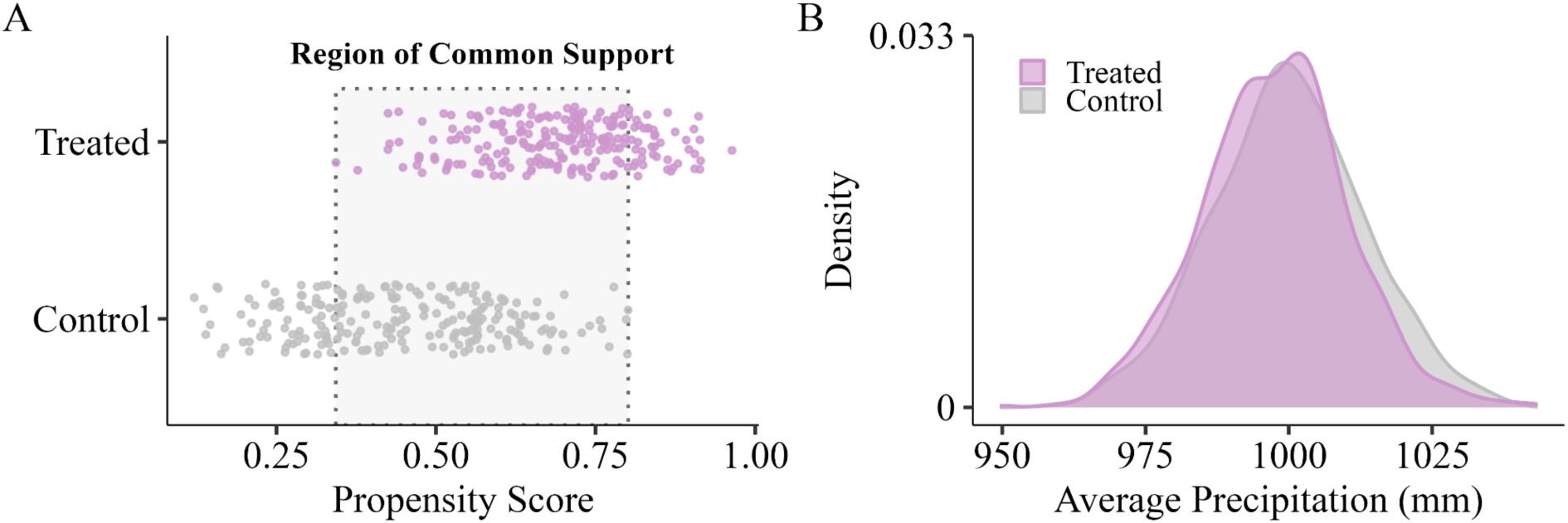
A) Region of common support (dotted grey box) determined for overlapping propensity scores within the treatment and control groups, B) Distribution of average precipitation for treatment and control groups within the region of common support.

### Matches control units on ecological rather than spatial proximity

Natural experiments often focus on spatial proximity between treatment and control groups, which may lead to confounding if the groups differ in site characteristics (Austin, 2011a). For example, control-impact (CI) studies compare treatment and control units sampled once after the treatment (Christie et al., 2019; De La Palma et al., 2018), often selecting controls within a specified distance from the treatment boundary (Mas, 2005; Andam et al., 2008). This design assumes that nearby units are directly comparable to treatment units because ecological conditions often become more similar with spatial proximity (Franco-Lopez et al., 2001; Nekola & White, 2004). However, spatial proximity alone may not account for fine-scale heterogeneity in confounding variables (Woo et al., 2021b). For example, nearby units may have similar climate characteristics (e.g. similar precipitation measured at > 800 m resolution), but differ in aspect, slope, and distance to road—variation that may confound the target causal effect (Woo et al., 2021b). Though spatial proximity is important to consider in plot designs, a more direct approach is to control for all confounders that would otherwise lead to systematic differences.

PSM addresses this challenge by focusing on both ecological and spatial proximity when comparing treatment and control units. Specifically, the propensity score (Step 4) is used to identify treatment and control groups within their **region of common support**—–i.e. the range for each identified covariate in which both treatment and control groups are comparable (*Figure 2*; Schleicher et al., 2020). Matching within the region of common support controls for confounding variables by only retaining ecologically comparable units. Geographic information can also be included in the region of common support to ensure close spatial proximity between each control and treatment pair (Papadogeorgou et al. 2019; Woo et al., 2021a; Woo et al., 2021b). For instance, Siegel et al. 2022a and Siegel et al. 2022b included latitude and longitude as covariates in the matching algorithm to minimize the spatial distance between each matched pair. If all confounding variables are identified and causal assumptions are met, the researcher can compare control and treatment units to estimate the treatment effect.

### Increases sampling efficiency

A major constraint in field ecology is the time, effort, and resources required to collect data, particularly in remote or difficult to access locations. As a result, researchers face tradeoffs among sample size, spatial coverage, and logistical feasibility (Field et al., 2005; Kemp et al., 2015). Classical field study designs often implement random sampling for population-level representativeness of important covariates in the regression (e.g. precipitation strata) (Steel et al., 2013; De La Palma et al., 2018). When field studies include many critical covariates, this can lead to an unfeasible sample size, or alternatively, the study results will inadequately account for all combinations of baseline covariates included in the regression model (Austin, 2011). PSM can increase sampling efficiency by identifying regions that are most important to sample (i.e. within the region of common support) to identify the causal effect (Stuart, 2010). Covariate combinations that do not fall into the target region will be dropped from the study, which can reduce sample size and increase sampling efficiency. While data collection within the region of common support focuses survey efforts, the estimated treatment effect may not apply to the dropped treatment units, thereby reducing generalizability of the study (see Considerations section below) (Crump et al., 2009).

## Case study: Quantifying wildfire effects on forest recovery

Here we describe a six-step framework that demonstrates how PSM can support causal inference from treatment effects in field ecology (*Figure 3*). These steps include: 1) identify the causal effect and outcome of interest, 2) identify confounding variables, 3) select the appropriate datasets and generate the unmatched dataset, 4) calculate the propensity score and select the matched dataset,

1. 5) evaluate the matched dataset, and 6) estimate the treatment effect. We demonstrate PSM using the US Forest Service’s Forest Inventory & Analysis (FIA) data, but the same steps apply when establishing new survey plots. Additionally, we highlight R packages, as well as research applications, that are useful for implementing PSM in the supplementary materials (*Supplementary Table 1*).

**Figure 3.**
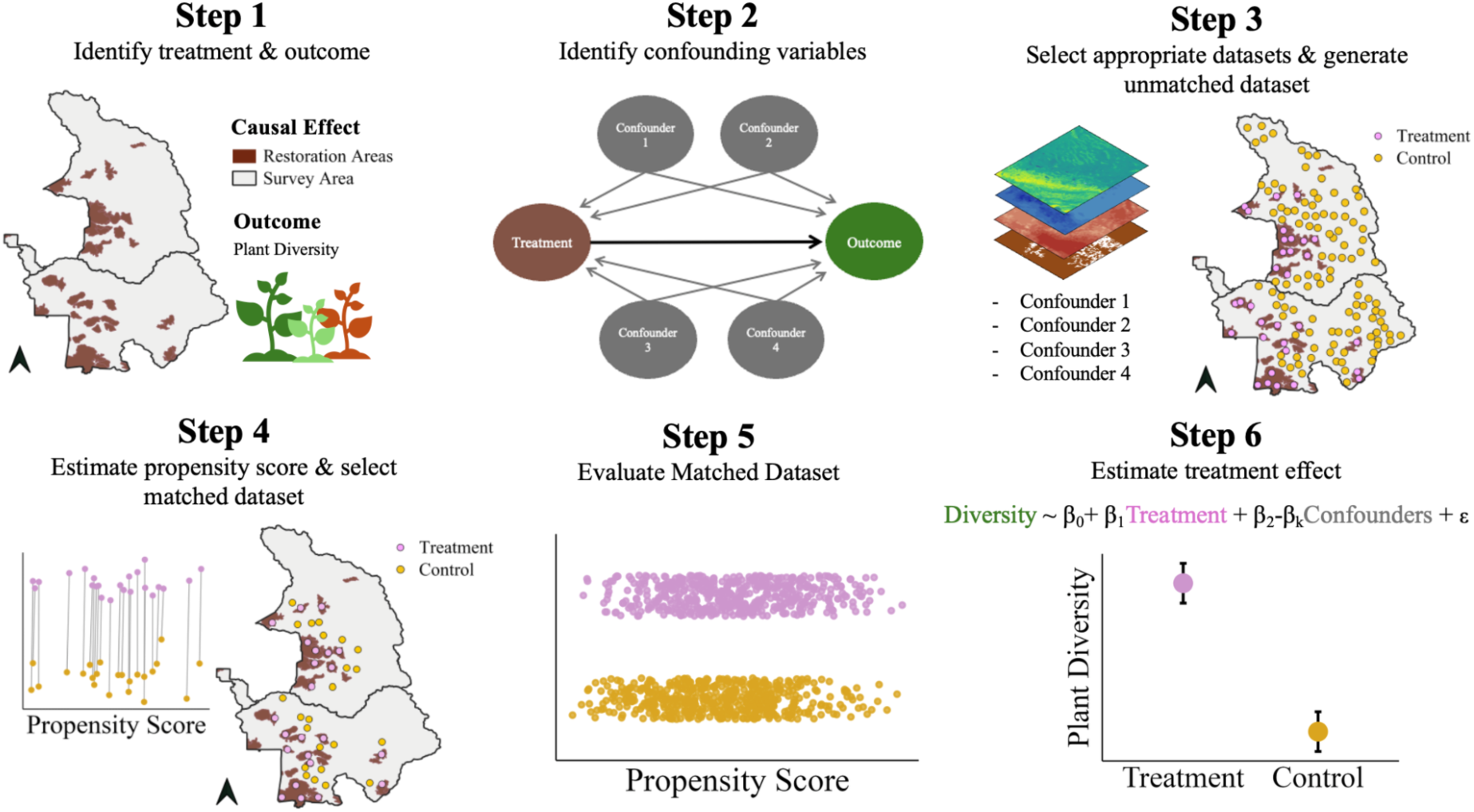
Six-step framework for implementing Propensity Score Matching. Step 1: Identify the target cause of interest (e.g., restoration treatments) and the outcome of interest (e.g., plant abundance). Step 2: Identify all confounding variables that could bias the estimated effect of interest. The figure displays a Directed Acyclic Graph that illustrates confounding variables using two arrows from the confounder to the treatment and the outcome. Step 3: Select the appropriate datasets for each confounding variable and generate the unmatched dataset. The figure shows spatial data for each confounder and randomly selected treatment (pink points) and control units (yellow points) that comprise the unmatched dataset. Step 4: Estimate the propensity score and generate the matched dataset. The figure shows the matched dataset, in which each treatment unit is paired with one control unit with the most similar propensity score. The map shows the distribution of treatment and control units from the matched dataset. Step 5: Evaluate the matched dataset visually and statistically. The figure visually evaluates the matched dataset using the region of common support, verifying that treatment (pink points) and control units (yellow points) have overlapping propensity scores. Step 6: Estimate the treatment effect. The figure shows the average difference in plant abundance between treatment and control units and the equation describes the regression model used to estimate the target causal effect (restoration treatments; pink) on the outcome of interest (plant diversity, green), controlling for confounding variables (grey variables).

### Step 1: Specify the target causal effect and outcome of interest

First, researchers carefully identify the target cause and the outcome of interest. The target cause is the specific variable whose causal effect is of interest. For example, the target cause could be a management intervention (e.g. prescribed fire, thinning), a disturbance event (e.g. bark beetle outbreak), or a continuous measure of disturbance severity (e.g. fire severity index). The outcome of interest is the variable that changes in response to the target cause—e.g. density, species richness, survival probability. Although it may be limiting to focus on one cause and one outcome (see Considerations section below), this step is critical to accurately identify confounding variables (Step 2).

In this case study we ask: “How does wildfire affect forest recovery in California?” The target cause is wildfire, defined as the occurrence of fire at an FIA plot between 1999-2018. We evaluate forest recovery using two plot-level outcomes: basal area and seedlings counts, which capture changes in overstory structure and post-fire regeneration (*Supplemental Methods S1*).

### Step 2: Identify all confounding variables

Next, researchers identify all potential confounding variables that could bias the estimated effect of interest (e.g. the effect of wildfire on forest recovery). Identifying confounding variables requires domain knowledge and careful consideration of ecological drivers within the system (*Figure 3;* Hernan et al., 2019; Arif & MacNeil, 2022). DAGs can be used to visually represent causal relationships and confounding variables by linking the treatment effect and outcome of interest with a directional arrow and including all relevant measured and unmeasured confounding variables, illustrated with arrows pointing to both the treatment and the outcome (*Figure 4*; *Supplementary Figure S1*; see Arif & MacNeil, 2022 for an overview of DAG construction).

In California, wildfire probability and forest recovery are jointly determined by many variables (*Supplementary Figure S1*). For example, hotter and drier regions are more prone to wildfire (Anderson-Teixeira et al., 2013) and are often associated with slower forest recovery (Davis et al., 2023). Wildfires occur more frequently at lower elevation on south facing slopes (Rogeau & Armstrong, 2017)—and both variables can slow forest recovery (Ireland & Petropoulos, 2015). Additionally, land-use history, such as management type and proximity to roads, influences the probability of burning (Siegel et al., 2022a; Mostafa et al., 2024) and forest recovery (Collins et al., 2017). Finally, forest type influences fire behavior and mechanisms for recovery (Anderson-Teixeira et al., 2013; Mallek et al., 2013). Our final **adjustment set** identified in the DAG (*Supplementary Figure S1*) includes the following confounders: topography, historical climate, land manager, average vegetation stress, pre-fire tree density, pre-fire vegetation type, distance to road, and management legacies.

### Step 3: Select appropriate datasets and generate the unmatched dataset

Once all confounders have been identified, researchers select appropriate datasets that measure each variable. There are four important considerations when selecting data. First, the data should match the spatial and temporal scales at which the confounding variable operates. Gridded climate data, for example, can range from 0.8–4 km resolution, which may be too coarse to capture microclimate variation if it is an important confounder. Second, researchers consider how well available data measures the confounding variable(s) of interest (Dudney et al., 2025a). For instance, existing datasets may provide an imperfect proxy (e.g. remotely sensed vegetation indices) for the confounding variable of interest (e.g. average vegetation stress), which may introduce bias. Third, researchers should assess the accuracy and reliability of the data. Measurement error can reduce the accuracy or precision of the estimated treatment effects and reduce the quality of the matched dataset (Hernán & Cole, 2009). Finally, researchers can filter the sample before matching if specific areas are influenced by processes that obscure the treatment of interest. For example, if the goal is to estimate ecosystem recovery following a disturbance, researchers may choose to exclude areas that were actively restored through plantings or other interventions to ensure the treatment is consistent across units.

For the case study, we utilize long-term data that measures post-fire forest recovery and ensure that measured variables match the spatial and temporal scale of each confounder. We then identify datasets with low measurement error and consider remote sensing metrics as a proxy for confounding variables that were not observed in the field. After collating the confounding datasets, we filter regions that are likely impacted by additional processes that could obscure the effect of fire on forest recovery. Here we use FIA data to causally identify the effect of wildfire on forest recovery in California (*see Supplemental Methods S1 for details about plot design and measured variables*) and compile additional data for variables that were not measured by the FIA program.

We first extract FIA plot-level data that measures topographic variables (slope, aspect, and elevation), distance to the nearest road, land manager (federal, state, or local) and geographic coordinates (*Supplemental Methods S1*). To compile confounding variables that are not measured by FIA, we extract 30-year climate normals (1991–2020) for precipitation and maximum temperature for each plot from PRISM at 800 m resolution (Daly et al., 2008). We use NDVI during the growing season (May–October) as a proxy for average vegetation stress leading up to the study period (1996–1998) (Landsat Collection 2 Tier 1 Level 2 32-Day NDVI Composite, U.S. Geological Survey) and ecoregions as a proxy for forest type (Griffith et al., 2016). Due to privacy concerns, plot coordinates are jittered by 0.8 km; we take extra precautions to spatially match climate and vegetation indices to each plot (*Supplemental Methods SI*). Finally, we control for potential confounding from historical management or disturbance by removing plots that experienced cutting, herbicide, tree planting or logging, as well as plots that burned multiple times, before and after the fire occurred. Tree density, a confounder identified in Step 2, was not measured prior to the study. Because PSM assumes no unmeasured confounding (Rosenbaum & Rubin, 1983), we can use sensitivity analyses to assess how robust estimated effects are to unmeasured confounding variables (see the Considerations section below).

Once the adjustment set has been measured and collated, control and treatment units can either be identified by the researcher or from existing data (e.g. FIA data)—comprising the unmatched dataset (Woo et al., 2021a). For example, Siegel et al. 2022a selected treatment and control units using an 800-meter grid placed across the study area. When treatment areas are unevenly distributed across space, researchers can randomly select units with minimum spacing between them, which can improve spatial coverage and reduce clustering. In this case study we used the California FIA database, specifically plots that were measured in 2019–2023 to capture an adequate sample size while capturing recent survey years. Within this subset of FIA plots, those that burned within 1999–2018 were added to the treatment group (n = 212), while plots that were not burned at any time were added to the control group (n = 857).

### Step 4: Estimate propensity scores and select the matched dataset

Using the unmatched dataset, researchers estimate a propensity score for each unit. The propensity score is the probability that a unit receives the treatment, controlling for confounding variables identified in Step 2. Propensity scores are commonly estimated using multivariate logistic regression, where treatment status is modeled as a binary response variable.

After estimating propensity scores, researchers identify comparable treatment and control units, using an appropriate matching method (Austin, 2010; Rosenbaum & Rubin, 1985). Common PSM algorithms include nearest neighbor and optimal matching (Rosenbaum, 2002), which pair treatment and control units with similar propensity scores until all control-treatment pairs have been identified or no suitable controls remain (Austin, 2010). Nearest neighbor matching selects the closest available control for each treatment unit, whereas optimal matching minimizes the total distance across all matched pairs (Austin, 2011; Stuart, 2010); the preferred approach depends on the research question and whether the final matched dataset achieves adequate covariate balance (Austin, 2011).

Researchers may determine the allowable “distance” between matched units using a caliper; a caliper of 0.2 or 0.25 standard deviations of the propensity score is common, though it’s appropriate value depends on the study design and balance diagnostics (Rosenbaum & Rubin, 1985; Austin, 2011b). Finally, matching can be performed with or without replacement—with replacement implies that a potential control unit can be used as a best match for more than one treatment. Matching with replacement may improve balance but requires additional analyses to account for control plots appearing in multiple pairs (Hill & Reiter, 2006).

Here we calculate the propensity score using the *MatchIt* package in R (Ho et al., 2011) as follows:

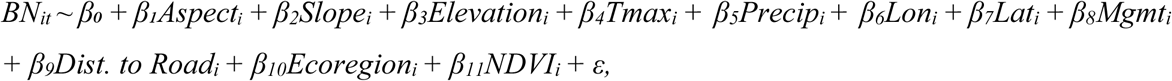

where *BN_it_* is the probability that a given plot burned, and the predictors represent confounding variables. We match without replacement, draw treatment plots from the unmatched dataset at random, select one control per pair, and apply a caliper of 0.25 standard deviations of the propensity score. We also use exact matching, which forces treatment and matched controls to have the exact same value for land manager, distance to road (categorical, S*upplementary Table 3*), and ecoregion. Unmatched plots are discarded, yielding a final dataset of 177 matched pairs (n = 354) across California. A researcher can use this same approach to identify sites for a field campaign, where the resulting matched dataset is a subset of the generated unmatched dataset established in Step 3.

### Step 5: Evaluate the Matched Dataset

Evaluating the quality of the matched dataset is a critical step because it informs whether matched plots are comparable—i.e. the plot selection has adequately controlled for confounding variables (Stuart, 2010). There are three important evaluation criteria that help researchers achieve the goal of maximizing similarity between control and treatment plots, while keeping as many treatment units in the unmatched dataset as possible (Austin, 2011; Schleicher et al., 2020). First, graphical diagnostics can evaluate similarity before and after matching. These include Q-Q plots and density curves, which compare the distribution for each confounding variable (covariate) between treatment and control groups in the matched and unmatched datasets, as well as plots of the propensity score distributions to help confirm the region of common support (Stuart, 2010). Second, researchers can evaluate whether treatment and control groups have similar means for each covariate using the **Absolute Standardized Mean Difference (ASMD)** (Rubin 2001; Austin, 2011; Stuart, 2010). ASMD estimates the mean difference between treatment and control groups for each covariate divided by the standard deviation, providing a unitless measure of balance (it is recommended that researchers aim for values close to 0 and no greater than 0.25) (Rubin, 2001). Finally, the variance ratio of propensity scores can be calculated by dividing the variance of the treatment group by the variance of the control group, capturing whether the two groups have comparable spread; values between 0.5 and 2 ensure comparable distributions between groups (Rubin, 2001). This complements the ASMD: treatment and control groups can have identical means but very different dispersions, meaning ASMD results may not always accurately evaluate similarity. Steps 4 and 5 can be iterated upon in the study design process until the quality of the matched dataset is sufficient for the research question.

For the case study, we follow the above-mentioned three steps to evaluate the matched dataset. Overall, treatment and control groups are well balanced across most observed confounding variables (*Table 1*; *Supplementary Methods S3; Table S4*). ASMD values are below the commonly used threshold of 0.1 for most covariates, suggesting that PSM reduced observed confounding bias. Exceptions include slope and two road categories, which had ASMD values of 0.12, 0.13, and 0.14, respectively, indicating modest residual imbalance. Variance ratios also suggest acceptable balance, with all covariate values ranging from 1.0 to 1.5 (S*upplementary Methods S3)*.

**Table 1.**
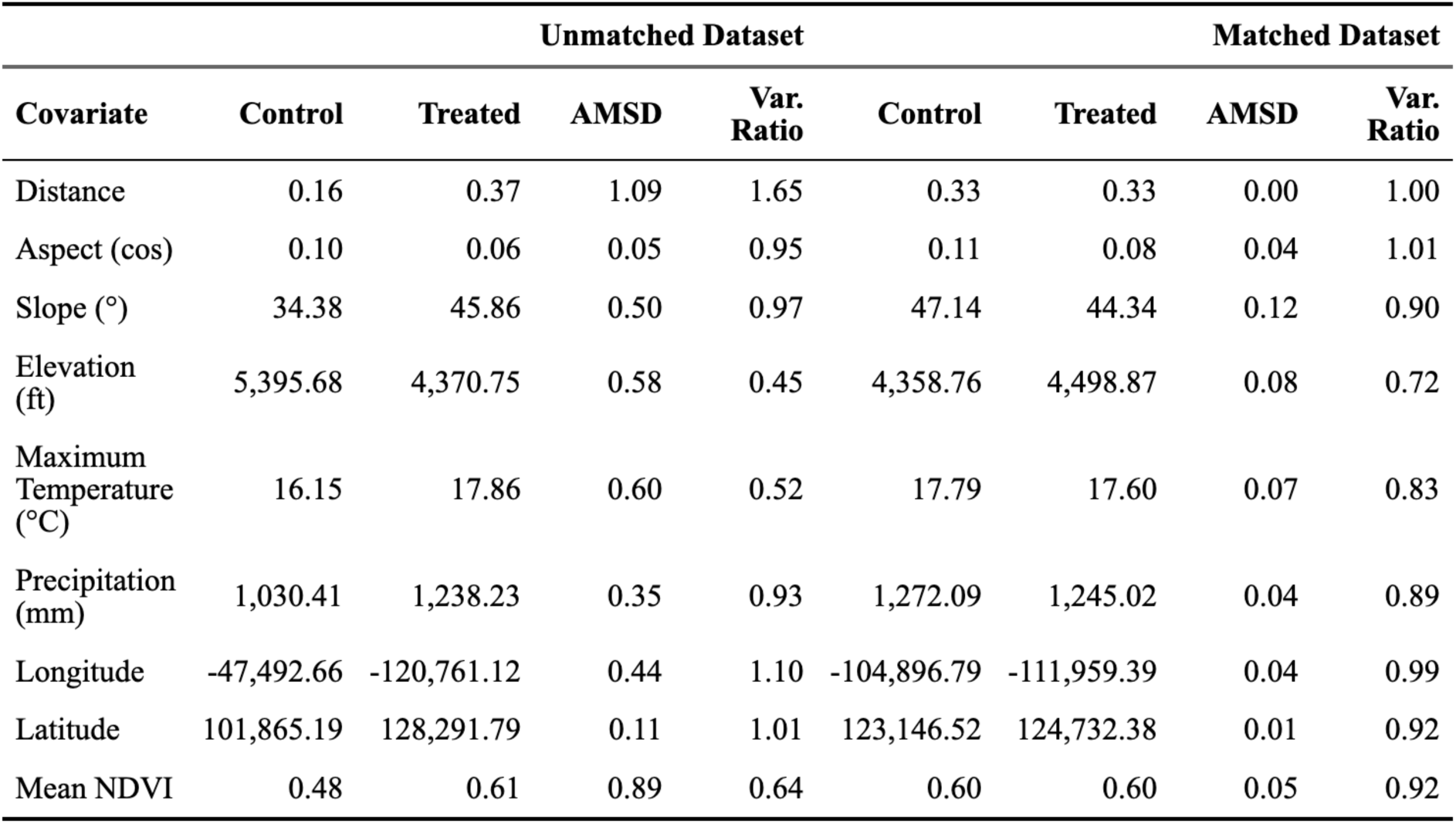
Differences in covariates, ASMD, and variance ratio (Var. Ratio) between the unmatched and matched datasets following PSM. Only a subset of covariates are included here for illustration purposes; the complete table of all covariates is included in the *Supplementary Materials Table S4*.

| Covariate | Unmatched Dataset |  |  |  | Matched Dataset |  |  |  |
| --- | --- | --- | --- | --- | --- | --- | --- | --- |
|  | Control | Treated | AMSD | Var. Ratio | Control | Treated | AMSD | Var. Ratio |
| Distance | 0.16 | 0.37 | 1.09 | 1.65 | 0.33 | 0.33 | 0.00 | 1.00 |
| Aspect (cos) | 0.10 | 0.06 | 0.05 | 0.95 | 0.11 | 0.08 | 0.04 | 1.01 |
| Slope (°) | 34.38 | 45.86 | 0.50 | 0.97 | 47.14 | 44.34 | 0.12 | 0.90 |
| Elevation (ft) | 5,395.68 | 4,370.75 | 0.58 | 0.45 | 4,358.76 | 4,498.87 | 0.08 | 0.72 |
| Maximum Temperature (°C) | 16.15 | 17.86 | 0.60 | 0.52 | 17.79 | 17.60 | 0.07 | 0.83 |
| Precipitation (mm) | 1,030.41 | 1,238.23 | 0.35 | 0.93 | 1,272.09 | 1,245.02 | 0.04 | 0.89 |
| Longitude | -47,492.66 | -120,761.12 | 0.44 | 1.10 | -104,896.79 | -111,959.39 | 0.04 | 0.99 |
| Latitude | 101,865.19 | 128,291.79 | 0.11 | 1.01 | 123,146.52 | 124,732.38 | 0.01 | 0.92 |
| Mean NDVI | 0.48 | 0.61 | 0.89 | 0.64 | 0.60 | 0.60 | 0.05 | 0.92 |

### Step 6: Estimate the treatment effect

Once adequate balance is achieved, researchers can sample control and treatment plots to measure the outcome variable. If the outcome variable has already been collected, researchers can move directly to statistical analysis. For example, a two-sample t-test can be used to compare the mean outcome between treatment and control groups (Austin, 2011; Andam et al., 2008; Welch, 1947). To improve precision, researchers can fit a regression model to estimate the causal effect of interest, using the same confounding variables previously used to estimate the propensity score (Imbens & Wooldridge 2009; Jones & Lewis, 2015). This approach can be particularly helpful if post-matching analysis suggests additional balance between treatment and control units is needed (e.g. because some plots were rejected in the field or were unable to be sampled due to time constraints).

We estimate the effect of wildfire on two forest metrics (basal area and seedling counts) using a fixed effect regression, implemented with the *fixest* package in R (Berge, 2018; R Core Team, 2025). To account for variation in fire years and the unequal distribution of the matched samples within each ecoregion, we included fire year and ecoregion as fixed effects. To control for variation in the timing of surveys following fire, we included the number of years between the fire and the FIA survey as a covariate in the model, assigning the same year to the matched controls. We found that wildfire has a significant effect on basal area (-63.51 ± 8.34, p < 0.001), but did not significantly affect seedling counts (1.05 ± 3.35, p = 0.75) (*Supplementary Methods S4*). Because we dropped treatment plots from our study, inference is limited to the treatment and control plots included in our region of common support.

Researchers can then assess whether their results are sensitive to potential unmeasured confounders by estimating bounds on treatment effects under different assumptions about the strength of unobserved confounding variables (i.e. Rosenbaum Bounds) (Rosenbaum et al., 2002). For example, because we were unable to include pre-fire tree density in the matching algorithm, we determined whether this unobserved confounder would influence the effect of fire on forest recovery. We estimated this effect using the *rbounds* package in R (Keele et al., 2010) to determine how the degree of departure from random assignment of treatment within a treatment and control pair could potentially bias the estimated effect (*Supplementary Methods S4*).

We computed two sensitivity tests that are included in Rosenbaum’s method for sensitivity analyses for matched data (*Supplementary Methods S5*; Rosenbaum, 2002). First, we used the p-value sensitivity test, which applies the Wilcoxon signed-rank test for each departure from random treatment (Gamma) (Wilcoxon, 1945). For instance, if the degree of departure were to equal two (Γ = 2), one unit would be twice as likely to receive the treatment, and hidden bias may influence the outcome (Rosenbaum, 2002). For Γ = 1–2.4, p-values remained < 0.05, which indicates that an unobserved confounder would need to increase the odds of wildfire by a factor of 2.4 to result in an insignificant effect on basal area. Second, we used the Hodges-Lehmann sensitivity test to calculate the bounds of the treatment effect by computing the median of each group given an increase in Γ (Rosenbaum, 2002; Hodges & Lehmann, 1963). Here we found that the bounds never crossed zero for Γ = 0–3; for Γ = 3 the bounds were 7.4–124.9, indicating that there is still a significant effect of wildfire even if one plot in a pair would be three times as likely to experience a wildfire given an unobserved confounder (*Supplementary Methods S5 & Table 7*). Therefore, we concluded that unobserved confounders likely do not bias the estimated effect of fire on forest recovery.

## Considerations When Implementing PSM

### Satisfying all assumptions required for causal inference is challenging

PSM rests on several assumptions that can be difficult to evaluate in practice. First, unbiased estimation depends on the assumption that there are no unmeasured confounding variables in the system (Rosenbaum & Rubin, 1983). In field ecology, this assumption is challenging to satisfy because critical confounders may be unknown or difficult to account for even after including all observed confounding variables (Siegel & Dee 2025). Because unmeasured confounding cannot be tested directly, sensitivity analyses can be used to evaluate how robust estimated effects are to hidden bias or identified, unmeasured confounders (Step 6; Cinelli & Hazlet, 2020; Stuart, 2010). For example, researchers can compare results across multiple control groups (Rosenbaum, 1987b), use placebo or falsification tests using outcomes or pretreatment covariates that should not be affected by the treatment (Imbens, 2004), or estimate bounds on treatment effects under different assumptions about the strength of unobserved confounding (Step 6; Rosenbaum, 2002).

SUTVA can also be challenging to satisfy, particularly in field ecology where discrete boundaries around treatments are relatively rare and disturbance events can shift the probability of subsequent disturbances (Ferraro et al., 2019; Dudney et al., 2025). Diffuse or poorly defined treatment boundaries can introduce hidden versions of the treatment, where control units may have received partial exposure (Schleicher et al., 2020). For example, nitrogen deposition from pollution can travel long distances via atmospheric transport and it can be difficult to spatially bound its effect (Wright et al., 2018). Where SUTVA is violated, researchers can remove affected areas, identify buffer zones near the treatment boundaries to reduce spillover effects, explicitly estimate spillover effects or identify regions where interference is assumed to be absent (Baylis et al., 2016; Andam et al., 2008).

### Suitable control areas can be limited

PSM is best implemented where there are many potential sample points, increasing the likelihood of finding control units comparable to treatment units while preserving adequate statistical power (Schafer & Kang, 2008). In practice, however, several challenges can shrink the number of comparable control units. First, violations of SUTVA (e.g., **spillover effects** from the treatment itself (Baylis et al., 2016)), can eliminate otherwise suitable control units. Fish populations in marine protected areas, for instance, may migrate (i.e. spillover) into adjacent unprotected habitat, which would be eliminated from the dataset to avoid violation of SUTVA (Schleicher et al., 2020). Second, land-use history can vary systematically across space, making it difficult to identify control plots that are both spatially proximate to treatments and comparable. Control plots that border a wildfire, for instance, may have been thinned preceding the fire; eliminating these areas reduces the number of control units and increases the spatial distance between treatment and control units. Third, disturbances—such as extensive wildfires, regional droughts, or widespread land-use conversion—can occur at very large spatial scales, where regions that escaped the disturbance may differ systematically in climate, topography, or vegetation. As a result, the remaining control sites may be poor **counterfactuals**, even after matching. When identifying suitable control areas, researchers often need to evaluate tradeoffs between reducing bias and preserving statistical power.

### Achieving balance across all covariates can limit generalizability

Identifying treatment and control units that are comparable across many covariates may limit generalizability of the study (Stuart, 2010). When treatments are strongly correlated with confounding variables, this may result in a narrow region of common support that removes treatment units from the study (Schleicher et al., 2020). For instance, in a study of wildfire effects on shrublands, burned units in extremely hot and dry locations could be excluded if no unburned control units occur in comparable climatic conditions. In some cases, dropping treatment plots could mean that PSM is unable to answer the target question. More commonly, dropping treatment plots reduces the generalizability of the study because inference is only drawn from plots within the region of common support—i.e., the effect of the treatment in extreme regions of a confounding variable(s) may not be estimable (Schleicher et al., 2020). Assessing the covariate balance before collecting field data, however, provides an opportunity to modify questions or evaluate tradeoffs between causal identification and generalizability.

### PSM estimates a single causal effect at a time

A key distinction of PSM is its commitment to specific treatment and outcome variables before analysis, which may constrain exploratory field studies where multiple drivers and responses are often examined simultaneously. For example, ecologists will collect a suite of measurements and include multiple predictors in a single model, sometimes interpreting all coefficients as meaningful effects (Dudney at al., 2020). In a PSM framework, however, covariates used to calculate the propensity score are control variables and are not to be interpreted causally—interpreting these coefficients as meaningful is referred to as the Table 2 fallacy (Westreich & Greenland, 2013). However, if multiple treatment variables are of interest and sufficient control and treatment units exist, separate matched analyses can be conducted for each effect of interest.

## Conclusion

As anthropogenic disturbances intensify, there is an increasing need for inferential frameworks that accurately identify causal effects to inform on-the-ground management. Although manipulative experiments remain the gold standard for causal inference, many anthropogenic stressors cannot be feasibly or ethically manipulated at relevant spatial and temporal scales. Observational studies are therefore essential to measure disturbance impacts on ecosystems. Commonly used field designs in ecology, however, do not make causal assumptions explicit or testable, making it difficult to infer causation in many settings. Here we describe how PSM can become an important addition to a field ecologists’ toolbox because it can often control for statistical bias that would otherwise confound estimated effects. Following the PSM framework will help ecologists build confidence in their research designs and identify causal effects, which has the potential to transform our ability to develop management strategies in the face of rapid environmental change.

## Acknowledgements

1. O. Ross is grateful to members and affiliates of the Landscapes of Change Lab for their feedback, including Michelle Mohr, Jenny Cribbs, Julie Edwards, Joe Celebrezze and Elijah McGill. We also thank the special issue for the opportunity to share this work. This article was prepared in part by employees of the U.S. government as part of official duties and is therefore in the public domain in the U.S. The findings and conclusions in this manuscript/publication are those of the author(s) and should not be construed to represent any official U.S. Government views or policy.

## Box 1. Glossary

**Adjustment set:** The variables identified in a DAG that must be controlled for (i.e., conditioned on)—through study design or statistical adjustment—to reduce bias and estimate the causal effect of interest.

**Causal Selection Bias:** Statistical bias that arises when ecosystem stressors or management interventions are not randomly distributed across a target population such that the factors influencing treatment exposure also affect the outcome of interest.

**Common Support Condition:** The requirement that the treatment and control units have sufficient overlap in their probability of receiving a treatment, given observed covariates. In other words, units must have a non-zero probability of being treated (Heinrich et al., 2010).

**Confounding Variable:** A variable that affects both the treatment (i.e., target causal effect) and the outcome of interest. If not accounted for, confounders can bias the estimated effect of interest.

**Control Units:** Sample units (e.g., plots, individuals, pixels) that did not experience the focal stressor or effect being studied.

**Counterfactual:** The outcome that would have been observed for the same unit under an alternative treatment condition—for example, the outcome of a treatment unit had it not been treated.

**Estimated Effect:** The estimated magnitude and direction of the relationship between an outcome of interest and a target treatment, stressor, or predictor.

**Propensity Score:** The probability that a unit receives the treatment, given its observed characteristics or covariates (Rosenbaum & Rubin, 1983).

**Region of Common Support:** The range of propensity scores where treatment and control units overlap sufficiently to enable valid comparison.

**Sample Selection Bias**: Bias that arises when sampled units are not representative of the target population (Boyd et al., 2023; Anderson, 2001).

**Spillover effect:** A violation of statistical independence in which the treatment applied to one unit affects the exposure or outcome of another unit.

**Statistical Bias:** Systematic error that causes an estimated effect to differ from the true effect. Bias can arise from non-random sample selection, measurement error, model misspecification, or other features of the study design or analysis.

**Stable Unit Treatment Value Assumption (SUTVA):** The assumption that each unit’s outcome depends only on its own treatment status and not on the treatment status of other units. SUTVA also assumes that there are no hidden or multiple versions of the treatment (Rosenbaum & Rubin, 1983).

**Treatment Units:** Sample units that experienced the focal treatment (effect) of interest.

